# Emergent Color Categorization in a Neural Network trained for Object Recognition

**DOI:** 10.1101/2021.06.28.450097

**Authors:** JP de Vries, A Akbarinia, A Flachot, KR Gegenfurtner

**Affiliations:** Experimental Psychology, Giessen University, Germany

## Abstract

Color is a prime example of categorical perception, yet it is unclear why and how color categories emerge. While prelinguistic infants and animals treat color categorically, several recent modeling endeavors have successfully utilized communicative concepts to predict color categories. Rather than modeling categories directly, we investigate the potential emergence of color categories as a result of acquiring visual skills. Specifically, whether color is represented categorically in a convolutional neural network (CNN) trained to recognize objects in natural images. Systematically training new output layers to the CNN for a color classification task, we find clear borders between new (non-training) colors that are largely invariant to the training colors. Using an evolutionary algorithm that relies on the principle of categorical perception we verify these border locations. These results provide strong evidence that color categorization emerges as a function of basic visual skills and provide a new basis for uncovering how they emerge.

## Introduction

Color vision is a prime example of categorical perception, and, as such, has received considerable attention across several research domains (Harnad, 1987). Being dependent on both linguistic and perceptual processing has made it difficult to pinpoint the mechanisms responsible for the emergence of color categories, and why particular colors are grouped the way they are. This has led to a protracted debate as to what extent categorization develops *universally* (independent of local language and culture) and to what extent it is *relative* to local communication (for an elaborate discussion on the Sapir-Whorf hypothesis see (Paul Kay, 2015; Paul Kay & Kempton, 1984). Proponents of the universalist view have pointed toward the overlap in focal colors across different cultures (Regier et al., 2005). Furthermore, it appears that categories can emerge independent of language development: Pre-linguistic infants pay more attention to color changes crossing categorical borders than color changes within categories ((Skelton, Catchpole, Abbott, Bosten, & Franklin, 2017) and several animal species respond to color categorically (Caves et al., 2018; Jones et al., 2001; Poralla & Neumeyer, 2006). Relativists (Davidoff, 2001), however, point to the difficulty children have acquiring color names (Roberson et al., 2004) and the case of a patient whose language impairments were associated with color sorting problems (Roberson et al., 1999). Also, while categorization has been found in birds and fish, researchers have failed to find categorization in some primates (e.g. baboons) and it has been argued that the methodology of other primate studies was biased towards finding categorical results (Fagot et al., 2006). Moreover, just as universalists point to the strong commonality among the development of color categories, proponents of the relativist view highlight differences (Roberson, Davidoff, & Davies, 2000; Roberson, Davidoff, Davies, & Shapiro, 2005).

With the apparent contradictions in findings, in more recent years, the universalist versus relativist debate evolved from contrasting two extremes, to looking at how different factors contribute to the process of categorization (P. Kay & Regier, 2006; Steels & Belpaeme, 2005). Importantly, recent advances have also started taking into account the varying utility of colors (Conway, Ratnasingam, Jara-Ettinger, Futrell, & Gibson, 2020; Gibson et al., 2017; Zaslavsky, Kemp, Tishby, & Regier, 2019). One seminal study shows that despite idiosyncrasies between cultures, overall, warm colors are communicated more efficiently than cool colors (Gibson et al., 2017a). Subsequent papers have demonstrated that utilizing a perceptually uniform color space in combination with concepts from communication theory, such as an information bottleneck (Zaslavsky et al., 2018) or rate distortion (Twomey et al., 2020) can be powerful in modeling the shape of color categories. While the degree of importance for communication in shaping categories varies, many of these recent studies rely on communicative concepts when it comes to shaping color categories. Notably, a recent study, where communicating deep neural networks played a discrimination game, demonstrated that allowing continuous message passing made the emergent system more complex and decreased its efficiency (Chaabouni et al., 2021).

While the modeling approaches incorporating communication principles have proven powerful in predicting categorization characteristics, the strong reliance on communication does not address the existence of what appears to be categorical behavior for color in the above-mentioned pre-linguistic infants and various animals. Also, a recent case study shows that color *naming* can be impaired while color *categorization* remains intact, emphasizing that in humans the link between communication and categorization can be decoupled (Siuda-Krzywicka et al., 2019). Furthermore, the trend at looking at the ecological relevance of color in shaping categories has also extended to animal research where it was found that host birds use a single-threshold decision rule in rejecting parasite eggs rather than the dissimilarity in color to their own eggs (Hanley et al., 2017) and female zebra finches categorically perceive the orange to red spectrum of male beak color (Caves et al., 2018). These latter findings, particularly, emphasize that while taking into account the variation in usefulness over colors through communicative concepts may be powerful, it does not preclude the possibility that the same usefulness variations can play a role during the acquisition of basic visual skill and, that this, by itself, could result in a categorical representation of color. Moreover, looking at the case study by Siuda-Krzywicka and colleagues it stands out that patient RDS tries to use objects to link names to colors (“this is the color of blood; it must be red” page 2473). A link between color categories and objects would be able to bridge the discrepancy between models that rely on communicative concepts to incorporate the varying usefulness of color, on the one hand, and the experimental findings laid out in this paragraph on the other. Moreover, finding that object recognition is a driver of color categories would be a crucial step towards finding the answer to <u>why</u> and <u>how</u> color categories spread across perceptual color space, the way that they do. The possible causes range from visual perception being shaped based on the physical properties of the world all the way to the need for efficient communication about color.

Focusing on color categorization as a potential emergent property of acquiring object recognition, here we investigate whether a categorical representation of color emerges in a Convolutional Neural Network (CNN) trained to perform an object recognition task on natural images. With this, our approach is not to model specific color category data directly, nor to model specific brain processes. Rather, we investigate whether a categorical representation of color can emerge as a side effect of acquiring a basic visual task; object recognition. The CNN is an excellent candidate for this purpose as unlike in any living species we can control its visual diet and train it on one specific task. Previously, many studies on the representation of color in a CNN rely on physiological style approaches (Engilberge, Collins, & Susstrunk, 2017; Flachot et al., 2020; Flachot & Gegenfurtner, 2018; Rafegas & Vanrell, 2018). Considering color categorization is likely a higher order process, we rely on classical principles from a long history of psychophysical studies to study the emergence of color categorization. The findings clearly show that a CNN trained for object recognition develops a categorical representation of color.

## Results

### Border Invariance

Perceptual research in non-human species requires indirect measures: in numerous species *match-to-sample* tasks have successfully been utilized to study visual perception for a long time (Kastak & Schusterman, 1994; Skinner, 1950). Training pigeons to match colors to one of three main color samples, subsequently allowed (Wright & Cumming, 1971) to introduce novel colors to evaluate to which of the sample colors they are matched and, consequently, find the points where pigeons switch from one color to another. Repeating the experiment with different training colors, they found crossover points to be similar across experiments, indicating a categorical perception of color. Here we use a similar approach to evaluate the color representation of a Resnet18 CNN (He et al., 2016) that has been trained on the ImageNet task, where objects in natural images have to be classified (Jia Deng et al., 2009). First, replacing the original output layer, a new classifier is trained on the network to classify stimuli containing a single word of a specific color (selected from narrow bands in the HSV color spectrum at maximum brightness and saturation). For stimulus examples see Figure 1A; the hue spectrum with corresponding training bands is found in 1B. In a second step we evaluate the retrained network using colors from the *whole* hue spectrum and determine to what classes these colors are generalized. As shown in Figure 1C, colors from outside of the training bands are largely classified to the neighboring color bands. As a consequence, the best color match for each point on the hue spectrum can be determined in a straightforward manner by simply taking the mode (see Figure 1D). The important question in regard to categorization now is what determines the transitions (or borders) between the found matches? One option is that the borders are dependent on the positions of the training bands, meaning a shift in these bands should translate to a shift in borders. Alternatively, the borders between the colors stem from a categorical representation of color in the network. In this latter case we expect the borders to be (at least partially) invariant to shifts in the training bands. To investigate this, we repeated the above process many times over while slightly shifting the training bands each iteration. As we do not know how many categories to expect, we vary the number of output classes from 4 through 9 (note that of the 11 basic color terms, only 6 are present in the selected hue spectrum). The result for 6 output classes has been visualized in Figure 1E: As the training bands are gradually shifted (indicated by the black lines) we find that the borders between categories appear largely invariant to these shifts in training bands.

**Figure 1.**
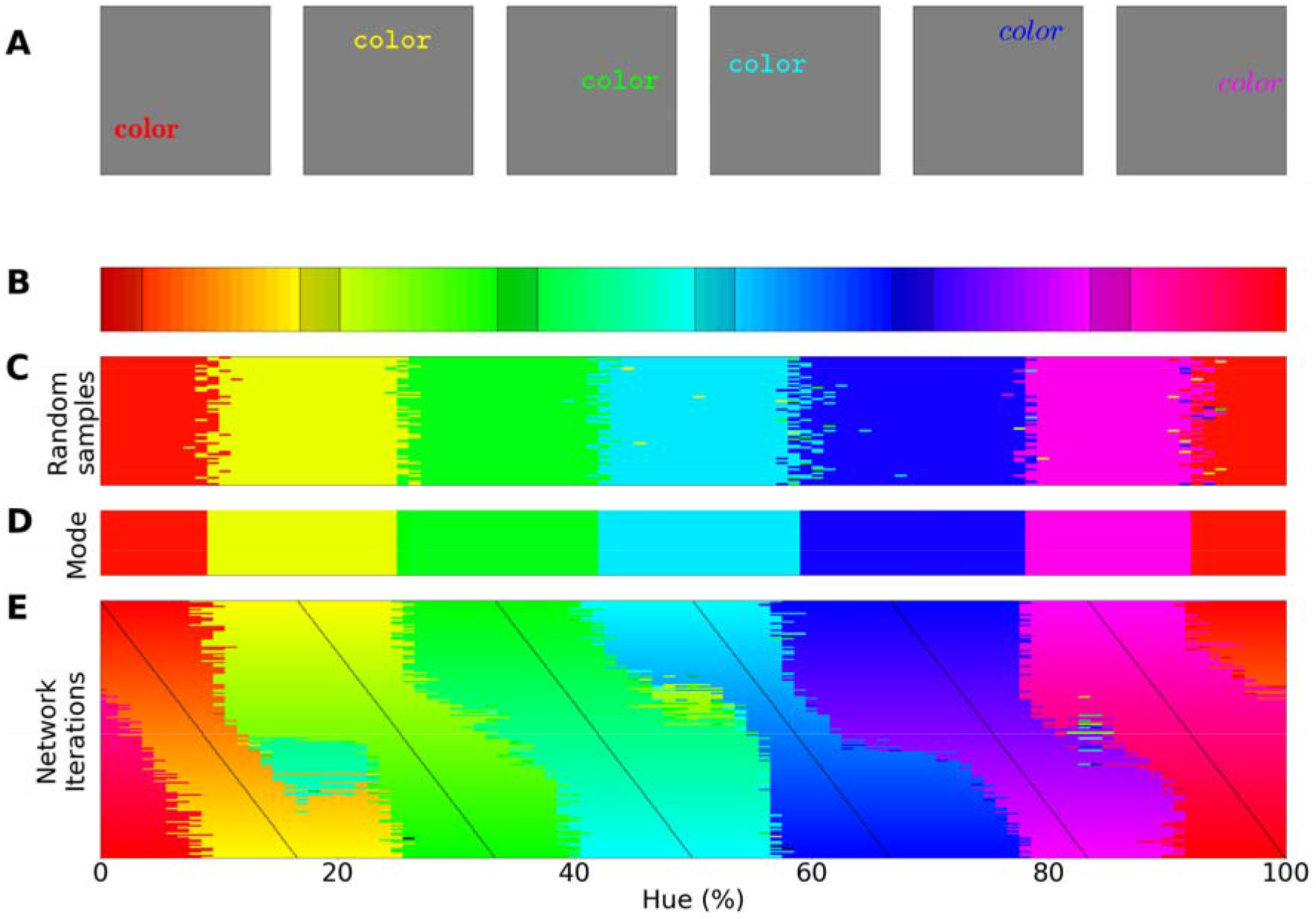
**A)** Six stimulus samples corresponding to the primary and secondary colors. **B)** Hue spectrum from HSV color space (at maximum brightness and saturation). The colors for each class are selected from narrow, uniformly distributed, bands over the hue spectrum. Bands are indicated by the transparent rectangles. **C)** Results from an individual training iteration for the bands depicted in **B**. In each iteration the same ImageNet-trained Resnet-18 is used, but a novel classifier is trained to perform the color classification task with the number of output nodes corresponding to the number of training bands. Each pixel represents a classified sample, colored for the class it has been assigned to (based on the hue of the center of the training band). **D)** A one-dimensional color signal produced by taking the mode of each column in **C**. In this manner we obtain an overall prediction for each point on the spectrum and can determine where the borders between classes occur. **E)** Results for networks trained on 6 bands on the hue spectrum. Each row represents the classification of a single network (as in **D**), trained on 6 bands, the center of which is marked by a black tick (appearing as black diagonal lines throughout the image).

To determine whether the borders are consistent across the shifting training bands, we plot the *transition count* that indicates the co-occurrence of borders in Figure 2A (i.e., the number of times a border occurs in a specific location, while the bands are shifted). Note that while Figure 1E only displays the results for a network trained with 6 output nodes, the full result depends on borders found when training the network with 4, 5, 6, 7, 8 or 9 output nodes. As those results are collapsed, inspecting the transition count shows that there are about 7 to 8 discontinuities in the hue circle. Using a simple peak detection algorithm that relies on neighboring data points to find peaks, 7 peaks are found (red dots in Figure 2A). Utilizing those peaks, we divide the hue spectrum in 7 different regions (Figure 2C), and averaging the colors in each region (weighing them based on the reciprocal of the transition count) results in 7 colors that can be seen as representing each category.

**Figure 2.**
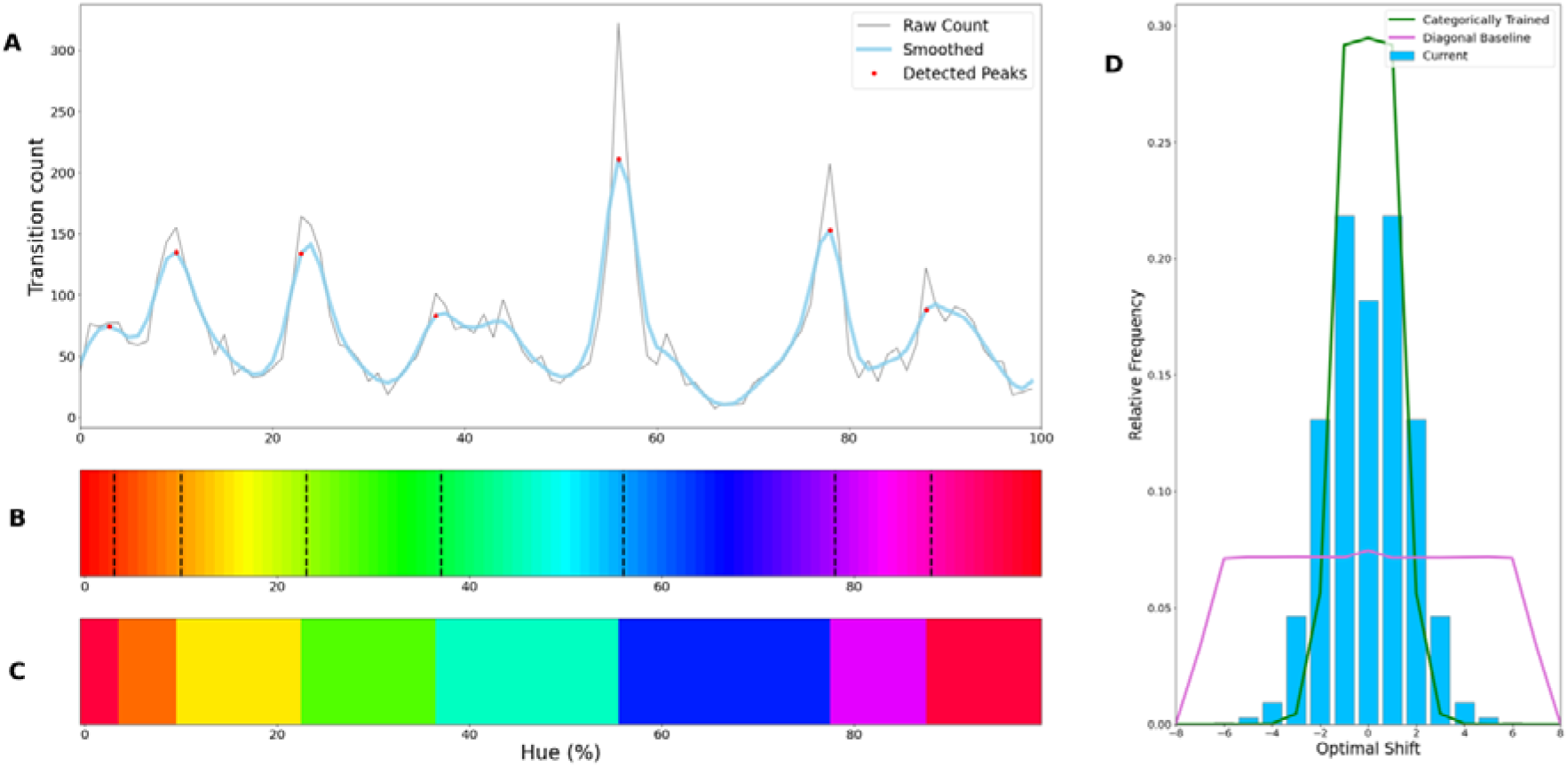
Border transitions in the color classifications. **A)** Summation of border transitions, calculated by counting the border transitions (as depicted in Figure 1D and E) for each point on the HSV hue spectrum (thin grey line). A smoothed signal (using a Gaussian kernel; blue thick line) is plotted to reduce the noise. Peaks in the signal (raw count) are found using a simple peak detection algorithm (findpeaks from the scipy.signal library) and indicated in red. **B**) The peaks superimposed on the hue spectrum as vertical black dotted lines. **C)** Category prototypes for each color class obtained by averaging the color in between the two borders (using reciprocal weighing of the raw transition count in **A. D)** For each row (as in Figure 1E) the optimal cross correlation is found by comparing each row to all the rows in the figure and shifting it to obtain the maximum correlation. In blue we plot the distribution of shifts for the when 7 output classes are used (as we appear to find 7 categories). For comparison we plot the result of a borderless situation (where borders shift with training bands) in purple and in green the result for a network trained from scratch on our 7 different color categories.

Considering the degree of invariance is not constant over the different borders, the question becomes what we consider sufficient evidence for finding categories and how we can exclude the possibility that this is a chance finding. To ensure the latter is not the case we have rerun the same experiment using similar architectures (notably a Resnet34, Resnet50 and Resnet101; see SI: Network Repetitions). In Figure S1 of the supplemental materials one can see, that given a similar architecture, similar border locations are obtained. Quantifying how “categorical” the current results are is more complicated, however. Naturally, if the color representation would be strictly categorical, the result in Figure 1E would primarily show straight lines. On the other hand, if the network incorporated some form a continuous representation of color, we would expect borders to be shifting with the shifting color bands and diagonal transitions, following the diagonal of the training bands, would be most prominent. In the first case, the highest cross correlation between individual rows would be found by keeping them in place as is. However, in the latter case, the maximum cross-correlation would be found by shifting the rows relative to each other. We can contrast these two cases by calculating, for pairs of rows, the lateral shift that produces the highest cross correlation and inspect the frequency for each of the found shifts. If the borders are stable across the different training iterations, we expect most shifts to be small (between -2 to 2). If the borders move along with the training colors, the shifts should be distributed more uniformly (−7 to 7; range is equal to the width of 7 uniform bands). Simulations for these two particular cases are shown by the green and purple curve in Figure 2E, respectively (see SI: Classification Simulation for more details). The histogram plotted in light blue shows the actual data, which more closely follow the green line, representing the categorical simulation. A Fisher’s Exact Test^1^ shows the difference between the count distribution is significant for all comparisons (p<0.0001; both pairwise and three-way). However, important to note is that the overlap between the current data and the shifting simulation is much smaller than with the categorical simulation (48% vs 74%). As such, despite a significant difference for both comparisons, it seems that the current findings more closely match with the categorical simulation. The significant difference between these two can be explained by the fact that directly training the color categories will lead to a less noisy color representation than when a network is trained on objects. Also, comparing the classification from the simulated data in Figure S2 (see SI: Simulation Classification) we can see that the color classifications from the current Resnet18 and the categorically trained Resnet18 are very similar and both deviate considerably from the continuous simulation.

We have elected to use the hue spectrum from the HSV color space as it includes multiple colors belonging to the basic color terms and because neighboring colors in HSV will be neighboring colors in the RGB space (in which our network was trained) at a constant distance. Varying the hue in the HSV space, however, does not only change the color of the word stimuli, but also their luminance. To ensure the current borders do not stem from a spurious correlation between color and luminance we have rerun the current experiment with different stimuli that include luminance distractors and a variable background luminance (see SI: Luminance Variation). While border locations are not a perfect one-to-one match with the current results, overall, they do seem similar and a categorical representation is again found. With the introduction of distractors and variation in background luminance the network can rely only on kernels coding purely for color and not a combination of color and luminance to perform the task. We also explored an alternative hue spectrum from a single plane of RGB color space. This led again to a categorical representation, albeit more noisy, presumably due to the reduction in chromatic contrast (see SI: Circular Color Spectrum).

### Evolutionary algorithm using principles of categorical perception

The generalization over neighboring colors and the invariant discontinuities between them are consistent with the notion that the network builds a categorical representation of color. Nevertheless, the lack of a broad understanding of CNNs in general and particularly their representation of color makes us wary to draw a definitive conclusion from only the first experiment. To evaluate whether the colors within the boundaries can indeed best be seen as belonging to categories we turn to the concept of *categorical perception*, where differences between colors within the same category are seen as smaller than differences between colors from different categories (Goldstone & Hendrickson, 2010). If the discontinuities we found indeed mark borders as in a categorical representation of color, we expect that generalizing colors falling between 2 discontinuities should be *easier* than generalizing colors that cross discontinuities. In humans, categorical perception is often studied using reaction time tasks (Paul Kay, Regier, Gilbert, & Ivry, 2009; Winawer et al., 2007; Witzel & Gegenfurtner, 2011), but a direct analogue of reaction times in neural networks is not available. There is however a different temporal performance measure available: the ability to evaluate how quickly a network can learn a task. If the boundaries found in the first experiment indeed resemble categorical borders, it should be faster for the network to learn to generalize colors within two neighboring discontinuities to a specific class than to generalize colors crossing discontinuities.

Specifically, in the current experiment we evaluate how well a set of borders fits the color representation of the network by evaluating how easily sets of 2 narrow training bands placed directly inside of each neighboring border pair can be generalized to single classes. One straightforward way of evaluating the discontinuities found above would be to make direct comparisons based on their locations. However, this would either mean using biased comparisons based on the borders found above, or require an almost infinite number of comparisons to ensure we compare the found borders to every alternative. To avoid this, rather than using the previously found borders as a starting point, we developed a search algorithm that uses the principle of categorical perception to find the optimal set of borders for the network from scratch. The only information taken into account from the previous experiment is that approximately 7 borders were found: An evolutionary algorithm is initialized with 100 sets of 7 randomly placed borders. Allowing the network to train for a limited number of epochs (the number of times the network sees every sample in the training set) we evaluate the fitness of each of the randomly initialized border sets. Using the *fitness* of each set, the best 10 performers are copied to the next generation (elitism) as well as a set of 90 novel border instantiations. The latter instantiations are generated by randomly selecting parents from the previous generation (with a bias for better performing ones) and recombining their borders to create new border sets. By allowing this process to run for some 40 generations it converges to a specific set of borders with little variation between the top ten performers. As evolutionary algorithms are not guaranteed to converge to a global optimum, we ran the algorithm 12 times to ensure the results are consistent. In the current case the borders should be less variable, but still line up with the peaks of the invariant border experiment.

Figure 3 shows where the evolutionary algorithm places the borders and we see a strong correspondence with the previously found borders (indicated by black vertical dotted lines). The only exception appears in the border between green and turquoise. However, note that this is also the region of color space where the transition count (as shown in 2A) did not show a sharp peak, as was the case for most other borders. Interestingly, this is also the border where the largest variability is observed for human observers (Hansen & Gegenfurtner, 2017). The current results suggest that the best explanation for the discontinuities we observed in the first experiment are that they indeed represent categorical borders dividing the color space. The fact that color categorization emerges in a CNN trained for object recognition underscores that color categorization may be important to recognizing elements in our visual world (Witzel & Gegenfurtner, 2018). This would also explain why there are strong universal tendencies in the development of color categories across cultures (Paul Kay & Regier, 2003). Further exploring the origin of the borders, the results shown in *SI: k-means clustering* indicate that efficiently representing the distribution of colors in the ImageNet database we used might, in part, explain the borders, in line with earlier results by (Yendrikhovskij, 2001).

**Figure 3.**
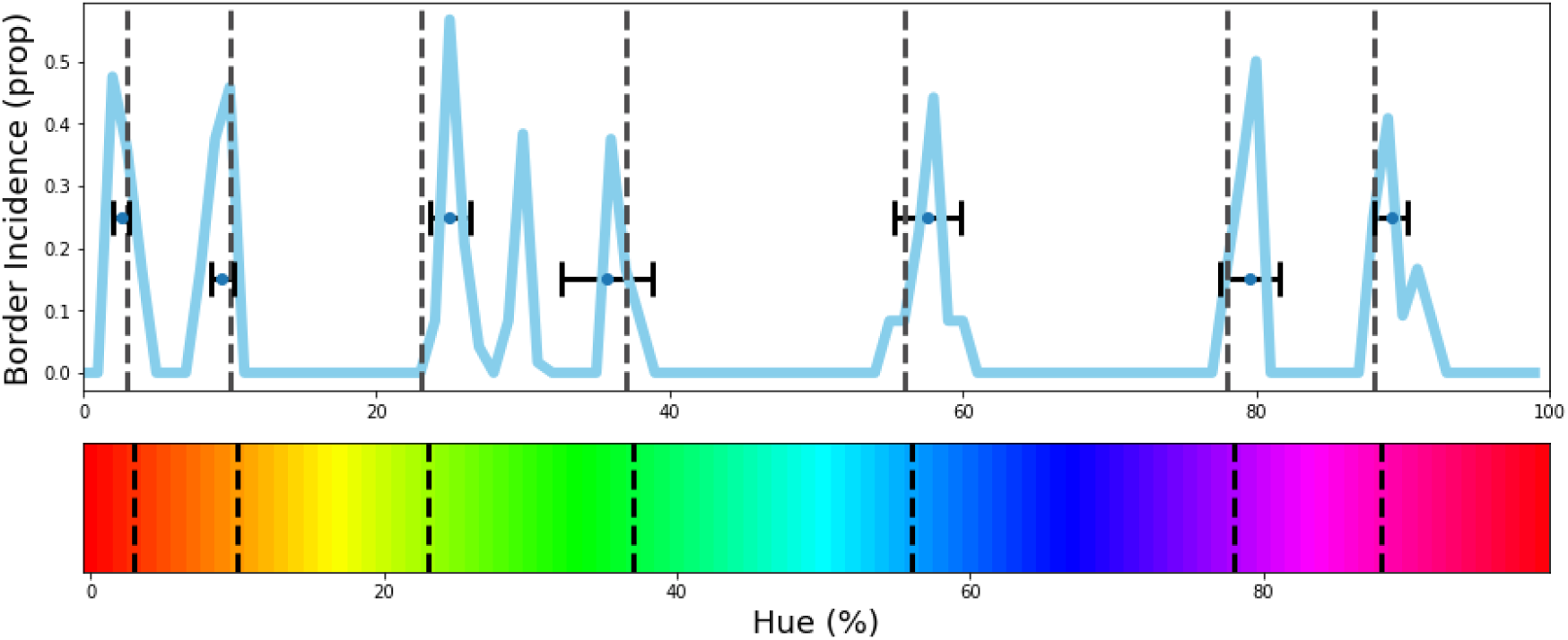
The evolutionary algorithm is repeated 12 times and we calculate the frequency of borders for in each of the top 10 border sets. The resulting frequencies are plotted in blue. Border-location estimates from the invariant border experiment are plotted in the graph and on the hue spectrum in dotted black vertical lines for comparison. Ordering the borders in each solution from left to right allows us to derive an estimate for each border the evolutionary algorithm produces by taking the median for the first, second etc. values, respectively. We have plotted these medians as points, including a horizontal errorbar that indicates a standard deviation to visualize the variability for these values. As can be seen, the estimate for each column agrees closely to the estimate from the invariant border experiment.

### Complex color stimuli

The converging results from the above experiments provide a strong indication that the network represents colors categorically. Still, the stimuli used above deviate considerably from the ImageNet database. We, therefore, wanted to ensure that the found borders are directly connected to how the network treats color. We proceed by investigating to what extent the borders generalize to tasks more similar to the one that the network was originally trained on. In a first step, we introduce a more complex set of stimuli that are comprised of multiple, colored, words on a randomly colored background to introduce contrast variations (see Figure 4A). Specifically, stimuli were comprised of 3 words colored based on the training specifications, as well as 2 additional words that were colored randomly. Finally, the background color is also randomly selected from the hue spectrum, however, at a lower brightness to ensure that there is sufficient contrast between words and background.

**Figure 4.**
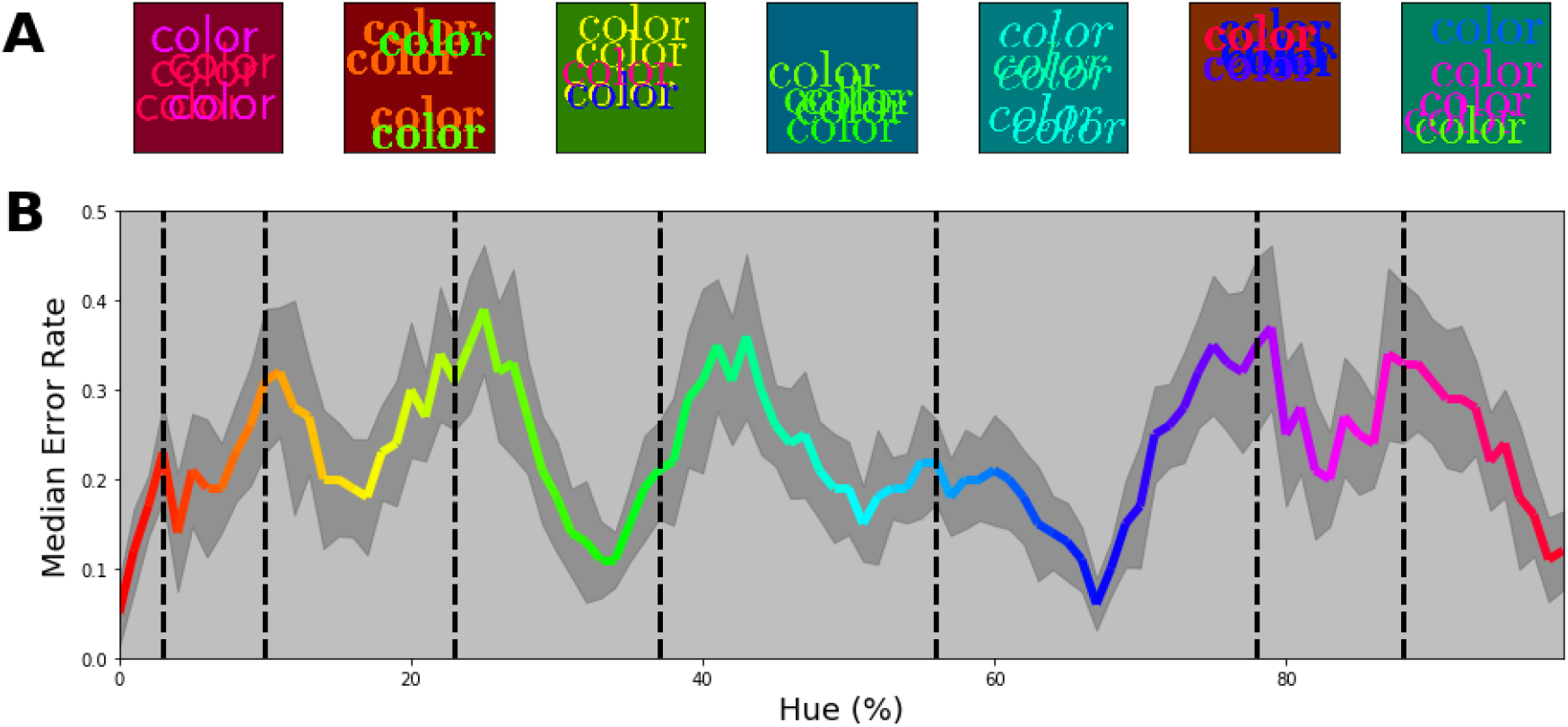
Multi-colored stimuli classification performance. **A)** 7 example stimuli, each sampled from a different color band. Each stimulus consists of 3 equally colored (target) words of which the color is determined by the selected class. Subsequently, 2 randomly colored (distractor) words, also randomly positioned, were drawn on top. These words are randomly positioned in the image. Finally, the background for each image is chosen randomly from the spectrum, but with a reduced brightness of 50%. **B)** Error proportion as a function of hue. Separate output layers have been trained on color bands that are shifted from the left to the right border in 10 steps for each category at the same time. This means, that while one network is trained to classify words of colors sampled from a narrow range on the left side of each category, another network is trained to classify words of colors sampled from the right side of each category for each respective class. After training the performance is measured using novel samples on the hue spectrum that match the color bands the network is trained is. Subsequently, the resulting error rate is displayed in the colored line by combining the performance for all the networks (shaded grey region represents one standard deviation). In this manner we can see the error rate typically increases as it approaches a border.

The key question is not whether the network can perform this task, but whether the categories obtained above are meaningful in light of more complex color images. Therefore, we trained novel output layers while iteratively shifting training bands from left to right within each category (in 10 steps). This allows us to evaluate performance as a function of the training band positions within the category. Two possible outcomes can be distinguished: On the one hand it is possible that for such complex stimuli the network deviates from the obtained categories and performance does not depend on where we select our training bands in the category. Alternatively, if the system can benefit from the categorical coding of color, we expect performance to be highest and the error rate to be lowest at the center of the categories, while the error rate should peak when the training bands align with the borders of the categories. In Figure 4B we see that this latter categorical representation is indeed what the network relies on: Overall, the network is able to perform the task reasonably well, but error rates are lowest towards the category centers, while increasing towards the borders. As in the experiments above, there is a slight divergence at border between green and turquoise.

### Recognizing colored objects

The previous experiment extends our finding to more complex color stimuli. We chose word-stimuli because they are made up of a rich set of patterns with many orientations. One notable element missing in these word stimuli are the large surface areas that are typically found in objects. In this experiment we investigated whether the previously found categorical borders still guide classification when classifying objects that incorporate large uniform areas. For generating objects with larger uniformly colored areas we rely on the *Google Doodle Dataset* (Ha & Eck, 2017). This dataset includes thousands of examples of hand drawn objects gathered from individual users. Because each drawing in the dataset is stored as a series of vectors it lends itself well to redraw the lines and fill the resulting shape with a uniform color (we plot some examples in Figure 5A). To further evaluate the usage of color categories in object recognition of the CNN we added one additional manipulation. So far, our experiments have aimed at looking on the reliance on color in isolation of potential other factors. However, with the introduction of objects of different shapes, a natural question is to what extent the network uses color or shape to classify the objects? To obtain a better insight into the interaction between these components we also raised the number of classes from 7 to 14. This allows us to evaluate whether the network simply ignores the color categories when they are not the sole source of discrimination, or can use them in combination with shape features.

**Figure 5.**
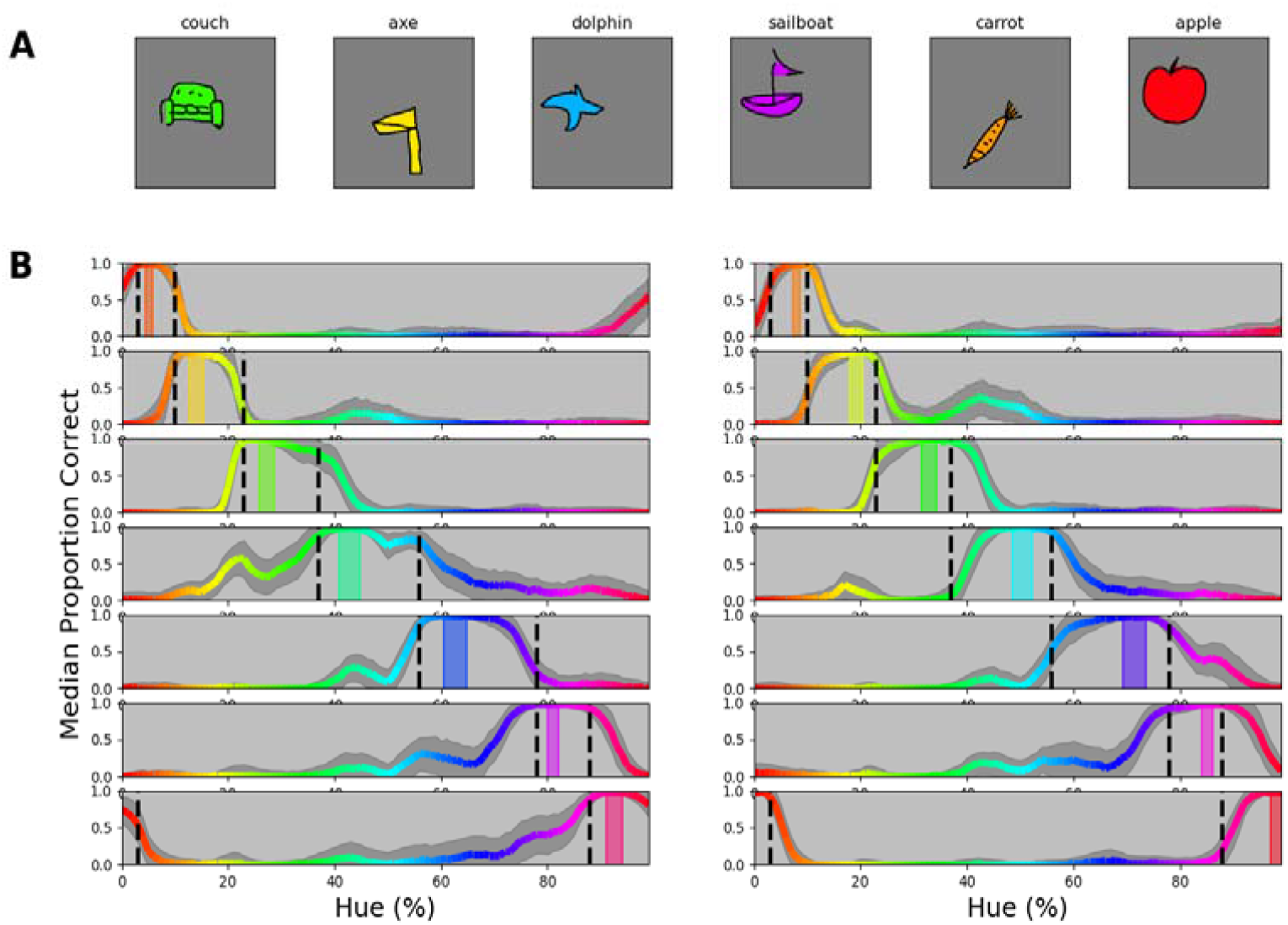
**A)** Samples of the google doodle dataset as colored by our simple coloring algorithm. **B)** Proportion correct as a function of hue. The 14 individual plots correspond to the 14 training bands that have been selected, 2 per category, 1 to the left of the category center, one to the right. The training bands are indicated by transparent rectangles on the spectrum, colored for the center of the band. The colored line represents the median proportion correct for the network on samples of the respective hue. Per instance, 80 drawings of the respective object are filled with each color on the hue (in 100 steps) and proportion correct has been calculated for each instance and hue individually. Lines correspond to the median proportion correct over the 100 repetitions (in each repetition the objects were randomly assigned to the various bands). The shaded grey area indicates the standard deviation over the 100 repetitions.

We ran several iterations, selecting random permutations of 14 objects for the 14 training bands on the HSV hue spectrum (2 per category). In Figure 5B we plot the results for each of the 14 bands, separately. Training bands are indicated using transparent, colored, bars, while the black dotted vertical lines again indicate category borders as obtained in the invariant border experiment. The proportion correct, plotted in the color of the hue for each object is measured over the whole hue spectrum (based on 100 permutations reassigning different objects to the colored bands); the grey shaded area indicates one standard deviation. We observe that performance is consistently high within the category borders in which the training band falls, while in most cases there is a steep drop in performance outside of the category bounds. This means that on the one hand the network uses color to distinguish objects from those that are colored for different categories. At the same time, however, it appears that to discern two objects within a category, classification relies on the object shape. As such, the network appears to combine both color and shape information and, important to the current research question, the representation of color it relies on, closely follows the previously found category borders.

## Discussion

The potential relation between cognitive concepts and language has sparked a lively debate (e.g. Casasanto, 2008) and, in this light, the potential relation between language and color categories has been extensively researched as a prime example of categorical perception. A number of recent studies on color categorization have focused on the development of categories by explicitly modeling their shape using communicative concepts. Here we have taken a different approach by evaluating whether color categorization could be a side effect of learning object recognition. Indeed, we found that color categorization is an emergent property of a Convolutional Neural Network (CNN) trained for object recognition. Importantly, our findings unite much of the previous research on color categorization. First, they can explain why the emergence of color categories over cultures broadly follows a universal pattern. Second, the findings are in line with the notion that color categorization emerges in pre-linguistic infants and animals. Third, the findings are in line with recent reports showing a dissociation between color naming and color categorization. Fourth, while the findings may seem in contradictory to recent models successfully utilizing communicative concepts to model category shapes, the varying communicative need these models rely on, likely correlates with the varying utility of colors to more basic visual tasks, particularly, those pertaining to objects. As such, the predictive power of these models may not rely on communication about colors per se, but on taking into account the varying utility over colors. Therefore, it is likely that the implicit incorporation of color patterns useful during object recognition leads to a categorical representation of color in a CNN.

Originally, color categorization was theorized to be culturally dependent (Ray, 1952). However, in 1969 Berlin and Kay proposed that color categories across a wide range of languages could be described using 11 basic categories, as such advocating that one universal process guides the development of categories across cultures. While this hypothesis was originally based solely on visual inspection of collected categorization data, later, evidence was provided by demonstrating statistical regularities across cultures (Paul Kay & Regier, 2003; Regier et al., 2005). The current findings can explain why the general development of categories is so similar across languages: If color categorization is a side effect of acquiring basic visual skills (given relatively similar circumstances across the globe) color categories are expected to shape in a similar fashion throughout many cultures. Naturally, this does not preclude local differences in the visual surroundings and communicative need from further shaping categories, however, the current findings do offer a potential cause for the broad similarities in color categories and their development.

Measuring the full hue circle in pre-linguistic infants, Skelton and colleagues (2017) found that the novelty preference was dependent on categorical crossings. As such it appears that some form of categorical representation develops early on. However, it is unclear whether these categories are the direct basis for categories in verbal communication. Four of the five categorical distinctions Skelton and colleagues found can be separated by the cardinal axis corresponding to the color representation in the retinogeniculate pathways, suggesting some of the categorical behavior may rely on the early representation of color (an issue pointed out by, e.g., (Lindsey et al., 2010; Witzel & Gegenfurtner, 2013). Also, color naming develops much later and children that have not yet acquired color names make recognition errors based on perceptual difference (Roberson et al., 2004). Similarly, it is unclear whether categorical representations of color in animals resemble those in humans. Nevertheless, recent findings have shown, that, as in humans, the utility of color may play an important role for the categorical color representation in animals (Caves et al., 2018; Hanley et al., 2017). As such, while the nature of the categorical representations in prelinguistic infants and animals is still somewhat unclear, the novel finding that a categorical perception emerges with the general acquisition of visual skills is in line with the current findings. Moreover, it is likely that any pre-linguistic categorical behavior lays the basis for subsequent color naming.

Similar to the approach in animals, we relied on a match-to-sample task for studying color categories in a CNN. With this indirect approach, some limitations are similar to those in studies on animals. Importantly, however, there are also clear advantages to studying categorization in a CNN. We were able to repeat the match-to-sample task for a great number of training colors (without the risk of inducing a sequential bias over training sessions) allowing for better estimates. Moreover, verifying the borders using the concept of categorical perception with an evolutionary algorithm is a computationally intensive task that cannot be straightforwardly applied to any living system, but is feasible for CNNs because of the speed at which an output layer can be retrained. Of course, compared to animal and infant models the CNN is the least convincing version of the adult visual system. Nevertheless, an important benefit is that as the access to activity of artificial neurons is complete it can be exploited in future research: While neural networks are often described as black boxes, compared to biological systems, artificial neural activity is much more accessible, as one can easily probe the activation of all neurons. With this kind of access, a logical next step is to investigate how the obtained categories are coded in the CNN. In humans, coding of colors seems to become narrower and more variable beyond the LGN (Kiper et al., 1997; Lennie et al., 1990) and colors belonging to the same category seem to be clustered together (Brouwer & Heeger, 2013). (Zaidi & Conway, 2019) suggested such narrowing may take place over areas (from V1 to IT) by combining earlier inputs (equivalent to a logical AND) or through clustering of local cells. CNNs can serve as a model for testing the viability of several of such concepts.

Where the terms color categorization and color naming are often used synonymously, the subtle distinction between them is key to the debate on the emergence of color categorization. From a strong universalist point of view, a color name is no more than a label applied to perceptually formed categories. From the relativist point of view, the direction is reversed and it is the existence of a color term that dictates the existence of the respective category (Jraissati, 2014). A recent case study shows a dissociation between color naming and categorization. Patient RDS is able to categorize colors, but his color naming ability is impaired (Siuda-Krzywicka et al., 2019, 2020). The dissociation between the two, favor a view where categorization is a process that can exist independently of the linguistical labels. Interestingly, despite the problems in color naming, the link between objects and colors was preserved (Siuda-Krzywicka et al., 2019). While it is possible to argue that our CNN does communicate (it assigns objects to a class), it is important to note that our base network at no point is required to communicate about colors directly, but at best about objects. As such, the current finding effectively lines up with the notion that colors and their categories may be formed as part of a way to identify objects.

The fact that the current categorical representation appears to emerge in the absence of color naming emphasizes that explicit color naming is not a necessity for the development of categories. This may seem to stand in contrast to many of the recent studies that use communicative concepts as a means to model the shape of categories (Chaabouni et al., 2021; Gibson et al., 2017b; Twomey et al., 2020; Zaslavsky et al., 2020). However, many of those studies derive their predictive power from combining these concepts with the non-uniformities in the utility across colors. As such, the communicative concepts could merely be a means to incorporate the varying utility across colors. For instance, where it has been demonstrated that warmer colors are communicated more efficiently than cooler colors (Gibson et al., 2017b), it has also been shown that objects are associated with warmer colors than backgrounds (Rosenthal et al., 2018). The latter emphasizes that the higher communicative need for warmer colors likely stems from their prevalence in objects. While we do not argue the process is devoid from communicative factors (it could be argued that the network communicates about objects), the current results can unify the previous findings by showing that acquiring a skill like object recognition can lead to the emergence of a categorical representation of color.

## Methods

### Invariant Border Experiment

#### Software architecture and Stimuli

The experiment utilizes a Resnet-18 as provided in the *models* module from the *torchvision* package (Marcel & Rodriguez, 2010). The network is initialized with the pretrained option on: weights are set based on having been trained on ImageNet (Jia Deng et al., 2009) a large database of human-labelled natural images. After initializing the network, we replace the output layer for object recognition with a smaller output layer with anywhere from 4 to 9 output nodes. The weights to this novel (replacement) classification layer are randomly initialized.

Stimulus images are generated using the *Pillow* package (Clark, 2015). Image size is the same as that used for the original ImageNet training (224×224 pixels). Each image contains the word “color” randomly positioned on a mid-grey background (font size 40, approximately 100 by 25 pixels, depending on font type). The color of the word is randomly (uniform) selected from the bands on the hue spectrum in HSV space. Color bands representing the individual classes are uniformly distributed over the hue space (see Figure 1A for stimuli sampled from example bands in Figure 1B). Brightness and saturation are set to the maximum level. The HSV color for each pixel is converted to its equivalent in RGB space as the network has been trained using three input channels for red, green and blue, respectively. The font of the word is selected from 5 different fonts.

#### Procedure

For each number of output nodes (4 through 9), we instantiate the network a 150 times, replace its output layer, and train each on a slightly shifted set of training bands. The combined width of the training bands equals 20% of the total hue range. During network training we only allow the weights of the novel classifier to be updated, the weights of all preceding layers remain as they were trained on ImageNet. Because we cannot determine the number of potential color categories a-priori, we vary the number of output classes from now 4 through 9. This results in training a newly initialized output layer for 150 (band shifts) times 6 (4 through 9 output classes) networks, making for a total of 900 training sessions. 500 samples are provided for each class and the network is trained for 5 epochs. During the training we keep track of the best network using 50 separate validation samples per class (from the same training bands). After training the network to classify the colors from the training bands, each network is evaluated over the whole hue spectrum by providing the network with 60 samples for each step on the HSV hue spectrum (divided into 100 steps). This results in 6000 classified samples for each of the 900 trained networks.

#### Analysis

The 6000 test samples for each trained output layer are used to determine the border crossings. In Figure 1C we plot the classification of these 6000 samples for a single training iteration. To determine the best prediction, for each step on the hue spectrum (each column in Figure 1C), we take the mode. In this manner we transition to a one-dimensional representation of the network’s performance on the evaluation task, with the prediction for each hue. Importantly, this one-dimensional representation, as plotted in Figure 1D, is used to determine the border crossings: for each network we determine the borders by simply picking the transition between predicted classes. Finally, we sum all the borders and from this we use a simple straightforward peak detection algorithm (findpeaks from the scipy.signal library) to find the locations on the spectrum where the borders are most invariant to change. To determine how “categorical” the found border invariances are, we determine the maximum cross-correlation for each row, compared to every other row, by shifting one of the rows and finding the optimal shift by looking for the maximum cross correlation. To ensure the circular nature of the hue is preserved hues are converted to 2D locations on a unit circle and a 2D cross correlation is run for the x and y coordinates. By obtaining the shift for each row, with all other rows, we obtain a distribution of shifts, that can be compared to distributions representing a categorical result and continuous color result. The former is generated by training a Resnet-18 (from scratch) on the currently obtained categories and, subsequently, evaluating it in the same manner as we evaluated the Resnet-18 trained on ImageNet. The latter distribution is determined by calculating the optimal shift for the case where the borders between colors move in parallel to the shifting bands.

### Evolutionary Experiment

Stimuli are the same as the previous experiment. We initialize 100 sets of 7 borders, each ordered from left to right (each representing a position on the hue spectrum). For each border set a Resnet-18 pre-trained on ImageNet is initiated and again the final fully connected layer is replaced. The stimuli for training the networks are generated via the same procedure as described in the Invariant Border Experiment, but now the hues for each class are selected from two narrow bands just inside of each set of neighboring borders (see Figure 6). Both band positions, as well as width, are relative to the two adjacent borders; The band starts slightly inside the border (with the closest edge located 5% from the border) and the band width is set to 10% of the total distance between the borders. We use two narrow bands at the ends of the potential category as this will mean that when the borders cross a categorical boundary the network will have to learn to generalize colors from different categories to single classes, while if the borders are located optimally, the 2 bands for each class will stem from 1 category each. We judge the fitness of a border set by evaluating how *fast* the network can learn the classes defined by the bands on the inside of two adjacent borders. Therefore, each network is trained for 3 epochs only, which is insufficient to reach peak performance, but allows us to evaluate which border set best fits the color representation of our Resnet: Border sets that align with the color representation of the network should allow the network to reach a higher performance quicker.

**Figure 6.**
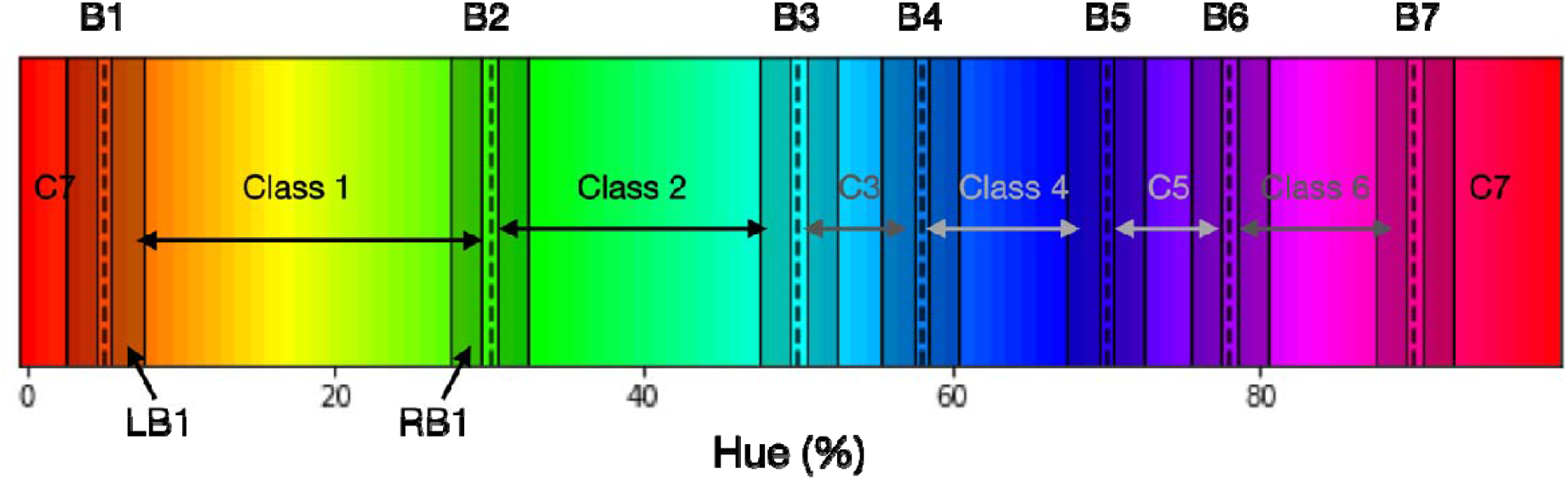
A single set of 7 borders (indicated by vertical dashed lines; labelled B1 through B7). Each space in between two adjacent borders represents a class. Colors for the training samples for, for example, Class 1 are randomly selected from one of the two bands, LB1 (Left Band for Class 1) and RB1 (Right Band for Class 1), on the inside of the borders of the class. Each of the two bands (not drawn to scale) comprises 10% of the space between; for class C1 this is the distance between first dotted vertical line (B1) and the second dotted vertical line (B2). Note that this means that the bands for class C3 for example are thinner than for Class 1. To ensure the training bands do not overlap at the borders, there is a gap (comprising 5% of the category space) between the border and the start of the band.

After training the 100 networks, the border sets are sorted by the performance of their respective network. Note that while we base the ordering on the performance of the network, the network is assumed a constant factor and the performance is attributed to the border set that is used to generate the training set; Essentially the network is the fitness function for the border sets. Once the fitness of each border representation has been established, the sets can be ordered by fitness and we generate a new generation of border sets. Firstly, through the principle of elitism, the top 10 performers of the current generation are copied directly into the next generation. Additionally, a novel 90 border sets that are created by recombining border sets from the current generation. Specifically, to create a new set, we select two parent border sets and, first, combine any borders that occupy similar positions across the two sets: This is done by averaging borders that are less than 5% apart between the sets (we start by combining the closest borders and the threshold is lowered when many borders are within a 5% range). Secondly, from the resulting borders, 7 are randomly selected to create a new border set. In order to converge to the optimal border-set, the selection of the two parent sets is biased towards the better performing border instantiations in the current generation as follows: 55% of the parents are selected from the 25 best performing border sets; 30% from the next 25 sets; and 15% of the borders are selected from the 25 sets thereafter. The bottom 25 border sets (in terms of performance) do not participate in the creation of offspring. To ensure some exploration occurs, in the offspring, some borders are randomly shifted. Specifically, we randomly select 2.5% of all borders and randomly shift them (random shift is normally distributed with an SD of 2.5% of the hue spectrum). The whole process is repeated 40 times. To allow for convergence after 30 generations random mutation is switched off.

### Multi-Color Experiment

#### Software Architecture & Stimulus

Again, the same Resnet18 trained on ImageNet is used as in the previous experiments. In the current version the ouput layer is replaced by one with 7 output classes, each matching a category. The stimuli were designed to include 2 factors that were absent previously. Firstly, we introduce multiple, colored elements in a single stimulus: Each stimulus class is still defined by a narrow color band on the hue spectrum, but we now draw 3 words into a 224-by-224 pixel image (we will refer to these as the *target* words). On top of that we add 2 additional words to the image of which the color is randomly selected from the hue spectrum (we will refer to these as the *distractor*s). All words are colored using the HSV hue spectrum at maximum brightness and saturation. Secondly, we introduce variations in color contrast. For the categorical borders to be meaningful in classifying stimuli their utility should not be overly dependent on color contrast (whether a banana is sitting on a green or a brown table, the network should still be able to use yellow to identify the banana). Therefore, we introduce a variation in color contrast by randomly selecting the color of the background from the hue spectrum. To prevent words from blending into the background we set the brightness of the background to 50%. Stimulus examples can be found in Figure 4A.

#### Procedure

Each network is trained on colors stemming from 7 bands, each based on the borders found in the invariant border experiment. The bands, making up 10% of each category range, are shifted from the left category border to the right border in 10 steps, for each step a novel classifier is trained 15 times to obtain a reliable average. After being trained on the borders we evaluate performance for each network: The error rate for the color bands networks were trained on is obtained by evaluating those networks for ticks on the hue spectrum that fall in that range. Subsequently, we plot the error rates for the different networks as a single-colored line (see Figure 4B), to demonstrate how performance varies, depending on training bands.

### Objects Experiment

#### Software architecture and Stimulus

Stimuli are generated using the line drawings available from the google doodle dataset (Ha & Eck, 2017). The database contains hundreds of objects, we selected a subset based on two criteria. First, we selected objects that lend itself to a simple color filling algorithm. This mainly consisted of finding objects that had clear outer borders with large spaces on the inside. Secondly, we prioritized objects that were reasonably consistent in regard to shape. The drawings are created by many different users, and, therefore, the approach could vary significantly. For instance, a user could have chosen to just draw only a cat’s head prominently, or include its entire body. Of course, such variations are not unlike the variation encountered in the images the network has originally been trained on. However, we are only retraining the output layer of the network. The latter means that the network has to rely on previously trained kernels to classify the shape, and a high degree of variation may not be easy to represent when only adapting the input weights to the last layer.

The drawing of the stimuli follows a simple procedure. Based on the drawn lines we create a color mask. This mask is created by determining for each pixel whether it is “enclosed” by lines. Enclosed, here, is defined by having a drawn line to its left (not necessarily directly adjacent), a draw line above it, to its right and below it. After the colored area is drawn, we draw, on top of it, the drawn lines at a thickness of 4 pixels. Example results of the process can be found in Figure 5A.

#### Procedure

The network is trained to classify a set of 14 objects (strawberry, apple, crab, dog, school bus, cow, dolphin, mushroom, bird, submarine, angel, sweater, sailboat, duck). For the training of each network, we selected 500 samples for each object from the google doodle dataset. We also selected another 50 samples per object to have a validation set to monitor the performance of the network throughout training. The fill color for all of these objects is randomly (uniformly) selected from a narrow band on the hue spectrum. Bands are selected to be non-overlapping and having 2 bands per category, one positioned right of the category center and the other left (each takes up 1/5 of the category bounds, one in the center of the right half and one in the center of the left half of the category). This results in 14 bands that are not always evenly spaced, or uniformly distributed throughout the spectrum. The bands can be observed in Figure 5B. To obtain a reliable estimate of performance on this task and to minimize the effects of the specific object selection we run the experiment 100 times, each with a different permutation of the object set with respect to the 14 color bands. After having trained the network, we evaluate the network using a separate set of 80 objects. Systematically changing the color of each of these 80 objects in 100 steps over the hue spectrum creates 8000 colored samples, that allow us to evaluate to what extent the object is classified based on its color.

## Acknowledgements

This work was funded by the Deutsche Forschungsgemeinschaft (DFG, German Research Foundation) – project number 222641018 – SFB/TRR 135 TP C2

## Footnotes

1 While the distribution in Figure 2D relies on comparing each individual row to all rows, this leads to the inclusion of doubles. For the statistical test we therefore only include each comparison once and take the shift as a positive distance value. The counts for each positive shift occurrence were entered into the test.

## Notes

### Competing Interest Statement

The authors have declared no competing interest.

### Summary of Updates

Figure 2D revised and text in Invariant Border experiment adapted regarding circular correlation analysis; Different ResNets have been tested and included in supplemental document; Simulations of Invariant Border Experiment have been visualized and included in the supplemental document; General text improvements

