## Supplemental Information for "Emergent Color Categorization in a Neural Network trained for Object Recognition"

**Network Repetitions (S1)**

To ensure that our interpretation of the peaks in the transition counts as invariant borders in the hue spectrum rather than noise is correct, we re-run the Invariant Border experiment a number of times. To introduce some further variation, we use the other Resnets available in the *models* module from the *torchvision* package. Specifically, we repeat the experiment with a Resnet34, a Resnet50 and a Resnet101, all of them pretrained on ImageNet. The results are plotted in Figure S1 below, Figure S1A repeats the results from the Resnet18 as presented in the main text for comparison. Comparing the transition count from the different networks (S1B-D) we clearly see a very similar pattern repeat. Most borders occur in approximately the same locations across the networks, only slight variations are seen as in some cases an extra border is found (around green and magenta) or a border is missing (between red and orange).

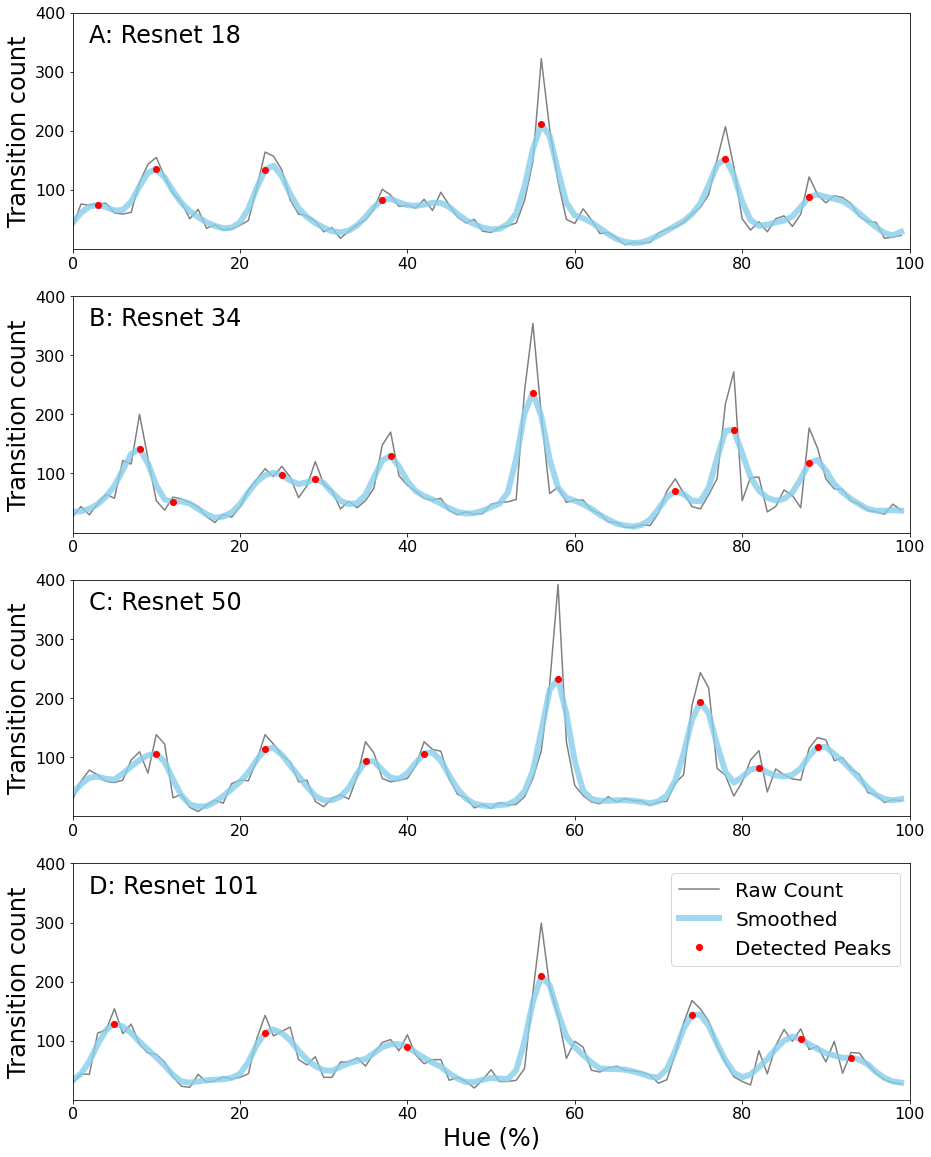

**Figure S1.** Transition counts for four different Resnets **A)** Transition count for the Resnet18 from the Border Invariance experiment of the main text. Transition counts are calculated by summing all transitions in the network’s color classifications going around the hue spectrum (see main document for more details). The thin grey line represents the raw counts, to remove noise we have also included a thicker light blue line which is a smoothed version of the raw count. Detected peaks are indicated using red dots. **B-D)** Same for the Resnet34, Resnet50 and Resnet101, respectively.

**Simulation Classification (S2)**

To determine to what extent the classifications of the Resnet18 on the color task can be seen as categorical we compare the classifications to two alternative cases. On the one hand we have the case where borders shift with the training bands, as could be expected when colors are represented in a (quasi) continuous manner. The shifting border simulation is plotted below in Figure S2 on the left. On the other hand, we have the case where the network relies on a true categorical representation. For this case we have trained a Resnet18 from scratch on the categorical borders found in the Invariant Border experiment and repeated the same analysis as in the Invariant Border experiment. The result can be seen in Figure S2 in the right plot. In the middle we have plotted the result from the original Invariant Border experiment for reference. As can be seen, the object-trained and the categorically-trained network produce very similar results, with the exception that the categorically trained result looks less noisy. A quantification of the differences can be found in Figure 2D of the article.

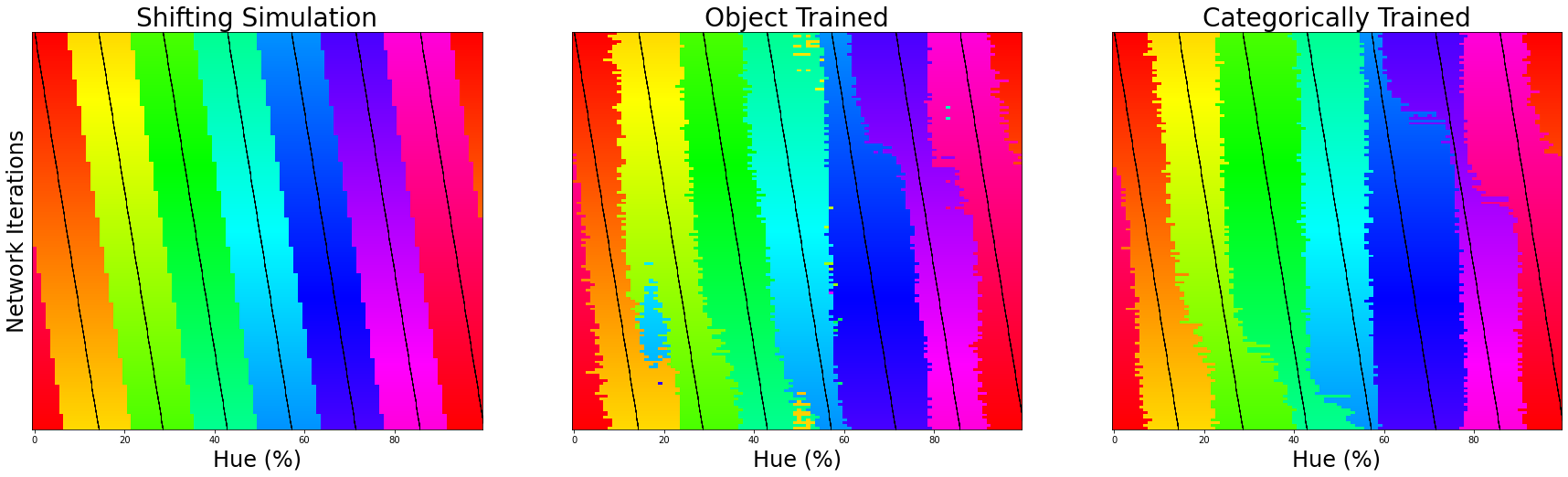

**Figure S2.** Classification simulation. We plot the color classification for the 3 cases from left to right (Shifting borders, data from the Invariant Border experiment and data from a categorically trained network, respectively). Each row represents the classification of a single network for 7 training bands, the center of which is marked by a black tick (appearing as black diagonal lines throughout the image).

**Luminance Variation (S3)**

In the invariant border experiment, we have drawn colored words on a uniform grey background. While varying the target word over the hue spectrum of the HSV color space, its luminance also changes. From this perspective it is possible that the obtained borders stem partially from a variation in luminance, rather than color. To ensure this is not the case, we repeated the same as experiment as before, but now with two important stimulus adaptations. Firstly, 10 greyscale words (random luminance) have been drawn into the background. Secondly, the background luminance is also randomly selected [range: 80-175 out of 0-255]. Stimulus examples are shown in Figure S3A. Otherwise the experiment is the same as the original.

The results are depicted in Figure S3B-D. While the locations of some borders are slightly shifted compared to the original borders, overall, we find a very similar pattern to those of the original experiment. As such, adding luminance variation does not change the notion that a categorical representation of color seems to underlie the current results.

Since the network can rely only on kernels coding purely for color and not a combination of color and luminance to perform the task, it is not surprising that the evaluation produces borders that are slightly different from those when luminance variations are not included.

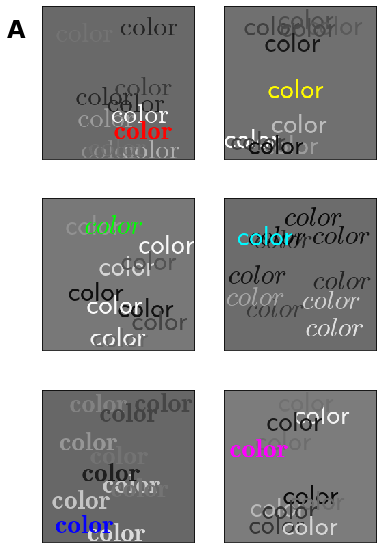

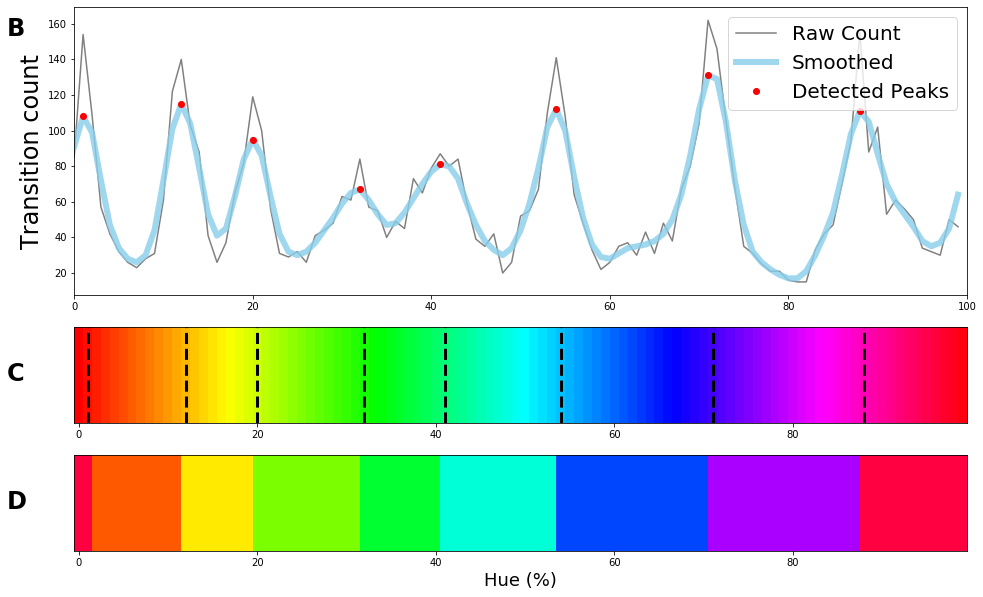

**Figure S3.** **A)** Stimulus examples drawn from 6 training bands aligning with the primary and secondary color. **B)** Raw transition count, i.e., the number of times borders between colors are found in a specific location as the network is trained on different training bands is plotted in grey. A smoothed version is plotted in light blue. The found peaks (based on the original count) are indicated in red. **C)** Found peaks indicated in the hue spectrum using black vertical dotted lines**. D)** Average color for each category. Average is weighted by reciprocal of raw transition count, making the troths in the data the most have weighted points.

**Circular Color Spectrum (S4)**

The hue spectrum from the HSV color space (at maximum brightness and saturation) follows the edge of the RGB color space (see Figure S4). To evaluate whether this irregular shape is a factor in the current results we repeat the experiment using a different hue spectrum. Specifically, we rerun the original experiment using colors picked from a single plane in RGB space, all at equal distance from the center (see Figure S4 for a contrast with the original hue spectrum). The novel hue spectrum is defined as a circle with a maximum radius (staying inside the RGB cube) in the plane of R+B+G=1.5. Repeating the original experiment (with a colored word on a uniform background) still shows a certain degree of categorical representation, but, looking at the raw signal count, the result is clearly noisier than the original experiment. Therefore, we have also repeated the experiment using the stimuli from Experiment S3 above. Adding luminance variation, we find a stronger correspondence between the borders from the HSV hue spectrum and the hue spectrum that follows a circular shape in RGB.

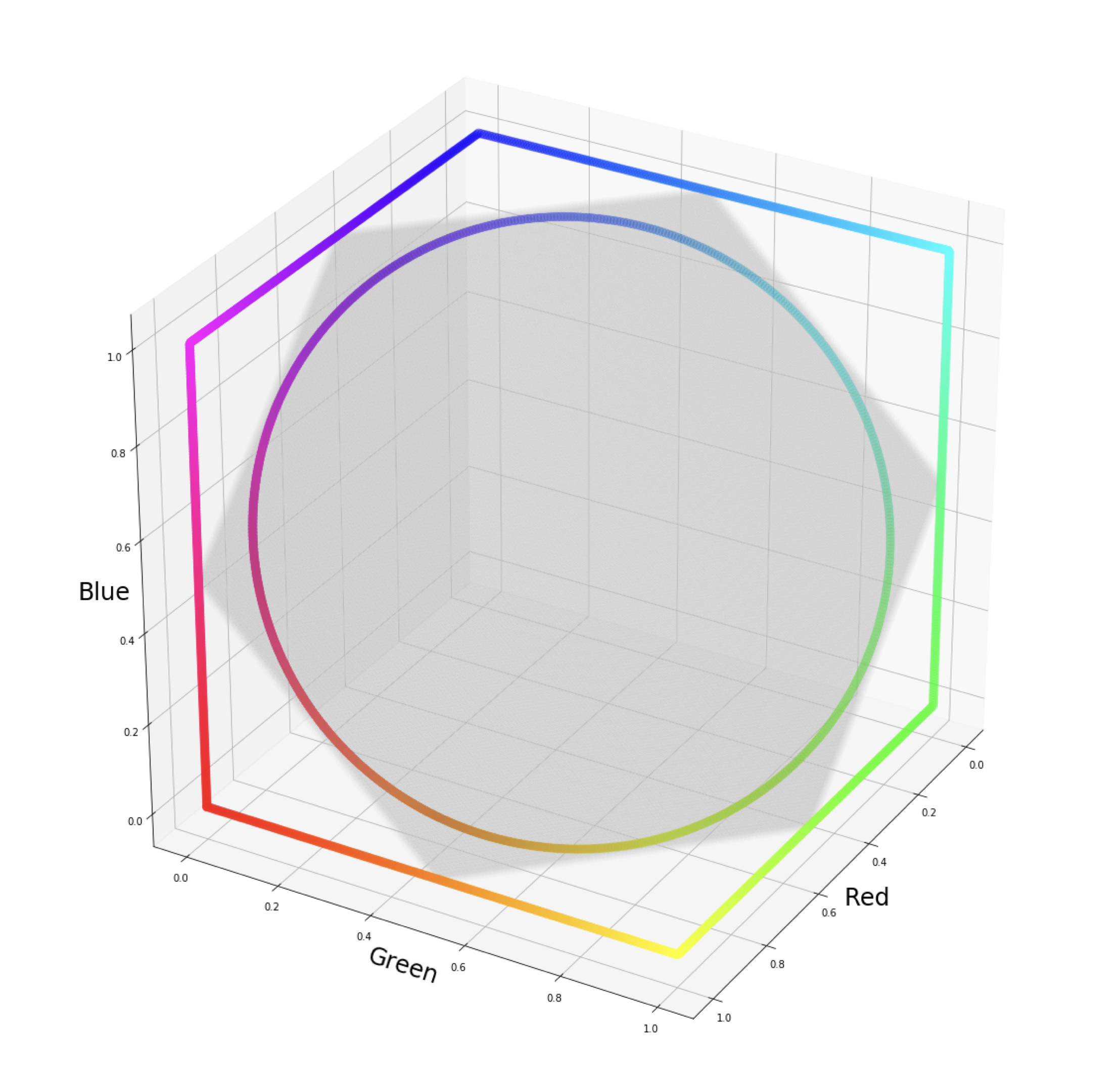

**Figure S4.** HSV hue spectrum and RGB hue spectrum displayed relative to RGB color cube. HSV hue spectrum at maximum brightness and saturation can be seen following the edges of the RGB color cube (subsection of RGB space in which values fall between 0 and 1). The RGB hue spectrum is defined as the maximum circle in the plane R+G+B=1.5. This plane is indicated in transparent grey.

Using single word stimuli from a hue spectrum following a circular shape through the RGB cube leads to a slightly noisier categorical representation (Figure S5A-C), presumably due to the reduction in chromatic contrast. Repeating the experiment with the stimuli containing luminance variation, however, does result in the same borders as with the HSV hue spectrum (Figure S5D-F). We hypothesize that the difference relies on the different kernels the network can rely on to perform the task. Likely, when getting closer to the center of the RGB cube there are even more kernels that classification can rely on. And, therefore, subtle luminance differences can be exploited and borders become less pronounced.

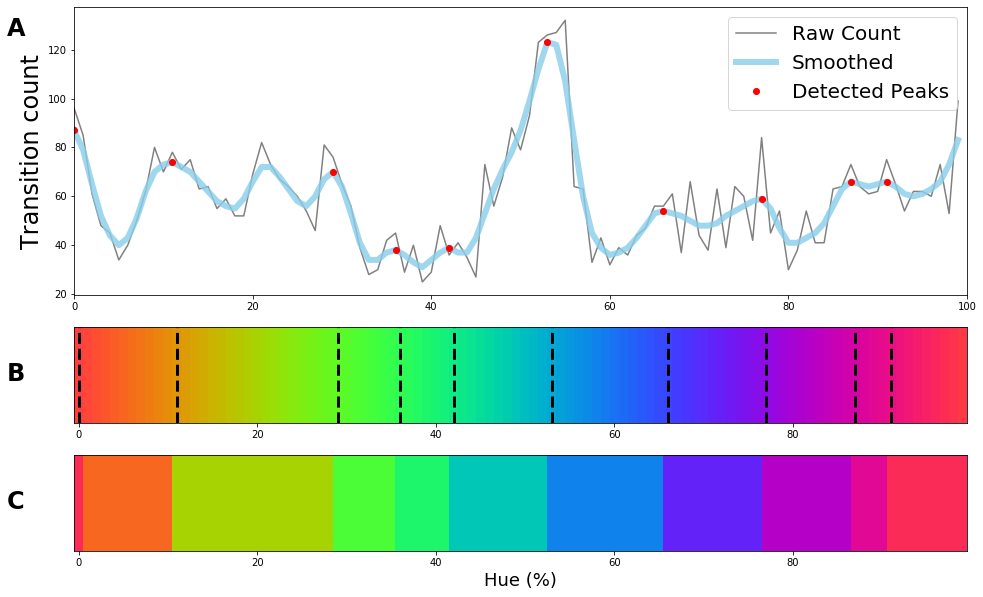

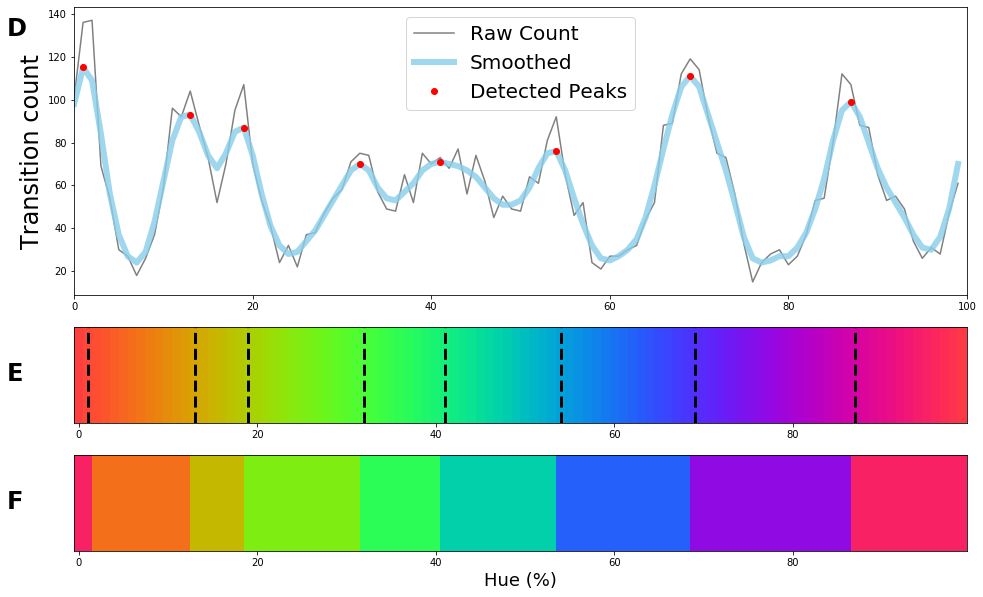

**Figure S5.** Left: Results from rerunning original experiment with stimuli sampled from custom RGB spectrum. Right: Results from rerunning the experiment with the stimuli defined in Experiment S1, but sampling colors from the custom RGB spectrum.

**Color clustering through K-Means (S5)**

Given the consistency of the borders over the experiments, the question arises what the origin of the locations of the found borders is. Previously, Yendrikhovskij (2001) demonstrated that performing kmeans clustering on colors from natural images results in cluster centers that are more similar to the focal color configuration than cluster centers obtained from uniform RGB values. To investigate whether the color distribution of pixels in the ImageNet dataset could explain the location the location of the current borders, we perform a kmeans clustering on them. For this we obtained the frequencies of colors in the HSV hue spectrum from a large sub-selection of images from the ImageNet database (subsection contained images that included bounding boxes; 535,497 images).

K-means clustering is performed by projecting the hues (taken as an angle 0-359) of the pixel values to the unit circle. The 2D coordinates are supplied to the k-means algorithm from the Scikit-learn package (Pedregosa et al., 2011) with a count of 7 for the *clusters* parameter. After clustering is performed the cluster means and cluster borders are projected back to the hue spectrum. In Figure S6 we see that there is correspondence between many of the borders obtained by the k-means algorithm and those found in the convolutional neural network. However, given that the borders do not line up exactly it also cannot completely explain the current border locations. Nevertheless, the correspondence makes it likely that the image statistics drive the location of the borders between the colors.

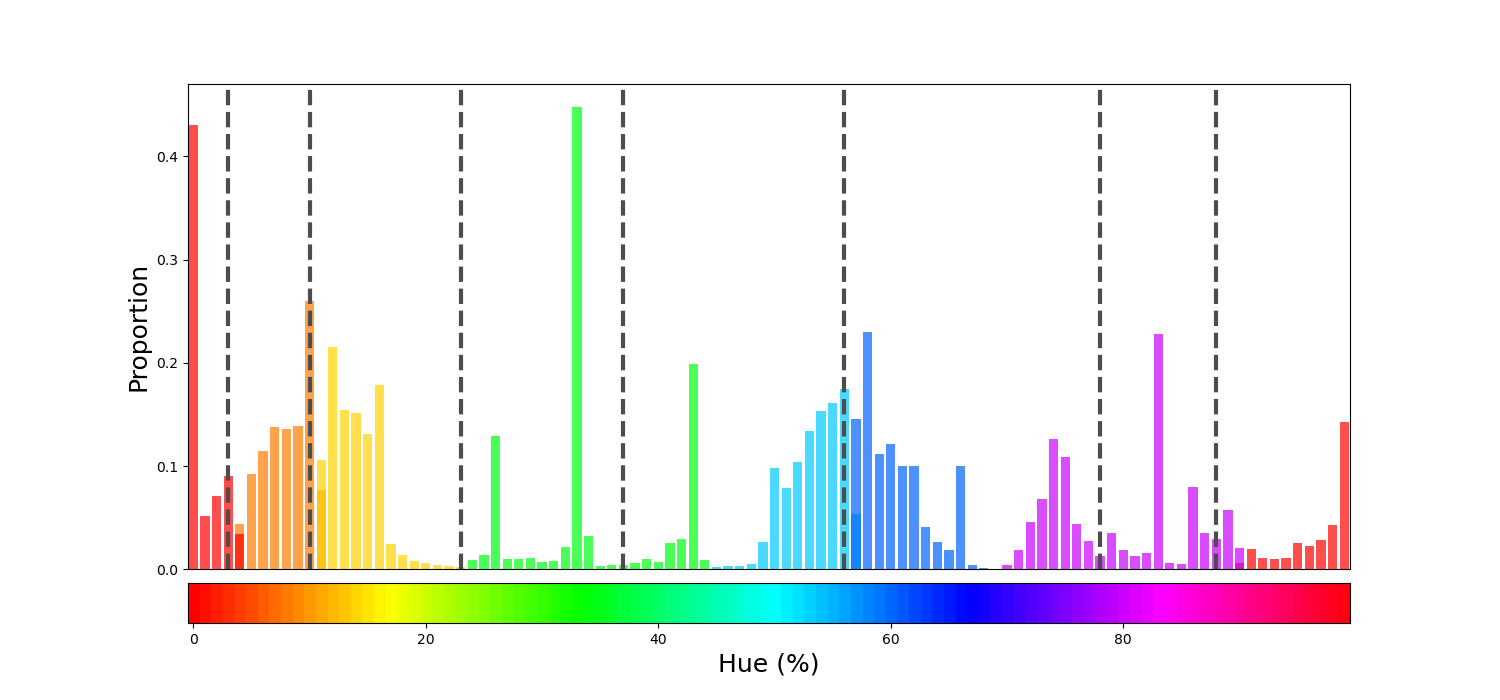

**Figure S6 Histogram of colors in ImageNet.** Colors from ImageNet have been selected to have a brightness and saturation exceeding 99%. The bars are colored in seven different colors, corresponding to the centers as found by the k-means clustering algorithm. As such, borders between clusters are indicated by color changes in the bars. The black vertical dotted bars indicate the borders as found in the original invariant border experiment.
